# Water-mediated interactions enable smooth substrate transport in a bacterial efflux pump

**DOI:** 10.1101/182683

**Authors:** Attilio Vittorio Vargiu, Venkata Krishnan Ramaswamy, Ivana Malvacio, Giuliano Malloci, Ulrich Kleinekatöfer, Paolo Ruggerone

**Affiliations:** Department of Physics, University of Cagliari, s.p. 8, Cittadella Universitaria, 09042 Monserrato (CA), Italy; Department of Physics & Earth Sciences, Jacobs University Bremen, Campus Ring 1, 28759 Bremen, Germany

## Abstract

Efflux pumps of the Resistance-Nodulation-cell Division superfamily confer multi-drug resistance to Gram-negative bacteria. AcrB of *Escherichia coli* is a paradigm model of these polyspecific transporters. The molecular determinants and the energetics of the functional rotation mechanism proposed for the export of substrates by this protein have not yet been unveiled. To this aim, we implemented an original protocol that allows mimicking substrate transport *in silico*. We show that the conformational changes occurring in AcrB enable the formation of a layer of structured waters on the surface of the substrate transport channel. This, in turn, allows for a fairly constant hydration of the substrate that facilitates its diffusion. Our findings reveal a new molecular mechanism of transport in polyspecific systems, whereby waters contribute by screening potentially strong substrate-protein interactions. The mechanistic understanding of a fundamental process related to multi-drug transport provided here could help rationalizing the behavior of other polyspecific systems.

---

Multi-Drug Resistant (MDR) pathogens represent one of the most pressing health concerns of the XXI Century due to their ability to elude the action of most (in some instances all) antibiotics (1–4). A special family of membrane transport proteins, the so-called efflux pumps, plays a major role in conferring MDR by shuttling a broad spectrum of chemically unrelated cytotoxic molecules out of bacteria (5–9). Polyspecificity and partial overlap among the substrate specificities of different pumps are striking properties of these efflux machineries (10, 11), making them a key survival tool of bacteria.

The efflux systems of the Resistance Nodulation-cell Division (RND) superfamily, which span the entire periplasm connecting the inner and the outer membranes, are mainly involved in the onset of MDR in Gram-negative bacteria (5, 12–14). The AcrABZ-TolC efflux pump of *Escherichia coli* is the paradigm model and the most studied RND efflux pump. It is composed of the outer membrane efflux duct TolC, the inner membrane-anchored adaptor protein AcrA, the small transmembrane protein AcrZ and the inner membrane RND protein AcrB (15). The lattermost is a drug/H+ antiporter fuelled by the proton gradient across the inner membrane and involved in the recognition and translocation of a very broad range of compounds (16).The multi-drug recognition capabilities and the postulated efflux mechanism of RND transporters are linked through an intriguing structural puzzle, which raised the question of how these proteins achieve their special features. An important step in this direction was made with the publication of the structure of AcrB (Figure 1A), revealing a asymmetric homotrimer in which monomers assumed different conformations, named Loose (L), Tight (T), and Open (O) [or, alternatively, Access (A), Binding (B), Extrusion (C)] (17–19). Such conformation was postulated to represent the active state of the transporter. A “functional rotation” mechanism was proposed explaining substrate export in terms of peristaltic motions induced within the internal channels of the transporter. In the simplest hypothesis (see e.g. (20–22) for a more complex picture), recognition of substrates should start at an affinity site, the Access Pocket (AP), in the L monomer (20, 23). Triggered by substrate binding, a conformational transition from L to T would then occur, accompanied by tight binding of the substrate within a deeper site, the so-called Deep or Distal Pocket (DP) (17–19). Successively, a second conformational change from T to O (supposed to be the energy-requiring step (24)) should drive the release of the substrate toward the upper (Funnel) domain through a putative exit Gate (hereafter, simply Gate (19)) (Figure 1B). After substrate release, the O conformation would relax back to L (coupled to proton release in the cytosol), restarting the cycle. Note that later different mechanisms of recognition were proposed for high vs. low molecular mass compounds, involving binding to the AP of monomer L and to the DP of monomer T, respectively (20).

**Figure 1.**
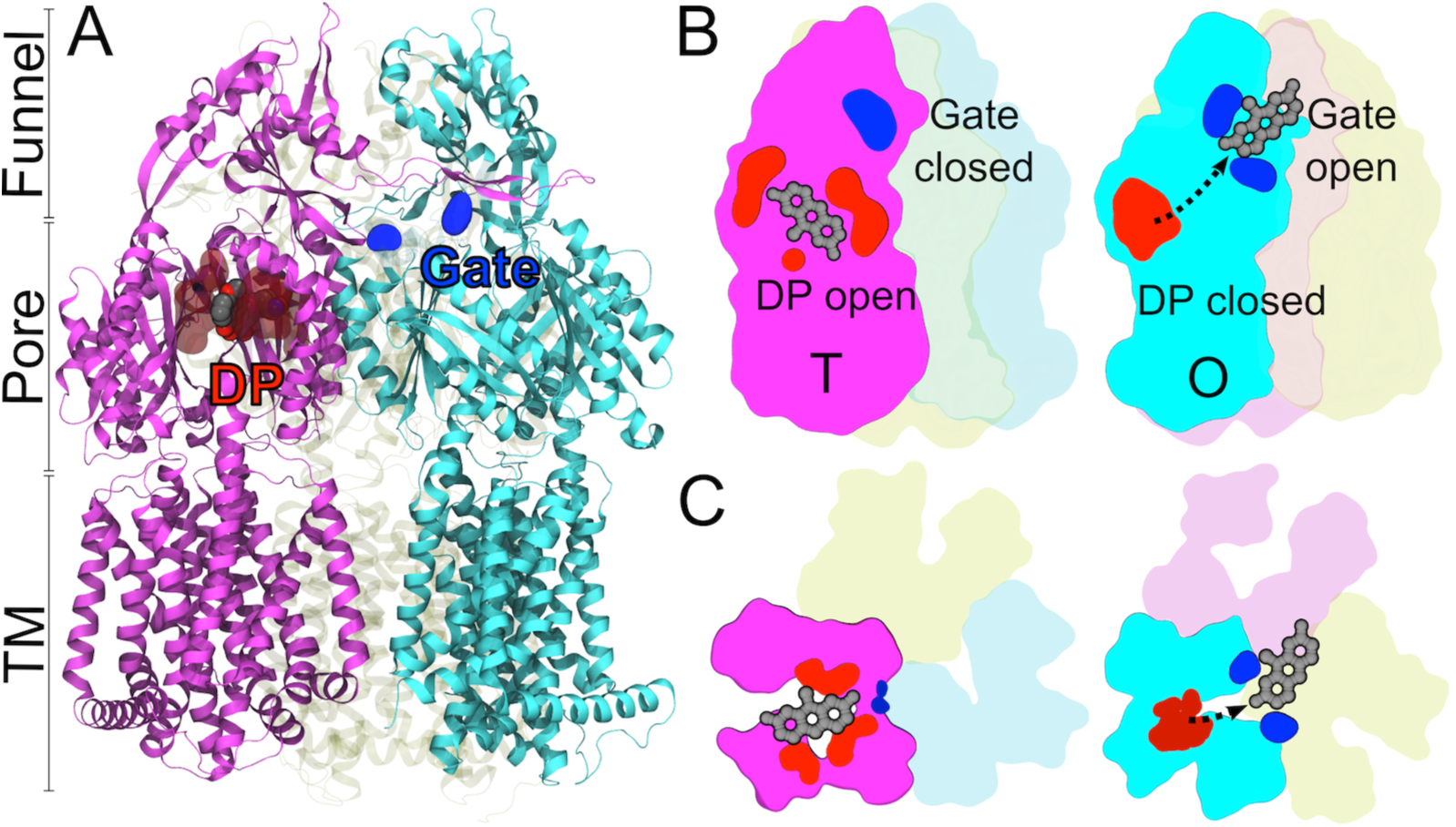
Structure of AcrB and schematic view of the T→O step of the functional rotation. (A) Overall structure of AcrB (PDB ID 4DX7 (23)). The three main domains TM, Pore and Funnel are indicated. Monomers are represented as ribbons, with T and O in front and colored respectively solid magenta and cyan, while L is transparent yellow. A molecule of DOX within the DP of monomer T is shown as spheres, with C, O and N atoms colored grey, red and blue respectively. The protein residues within 3.5 Å from DOX are shown as transparent red surfaces. Residues Q124 and Y758 lining the Gate of monomer O are shown as blue solid surfaces. (B) Schematic view of the T→O step of the functional rotation mechanism in AcrB (color code as in A). Left and right pictures represent L<u>T</u>O and T<u>O</u>L conformations respectively. The underlined monomers contain the substrate being transported, and are shown as solid shapes in contrast to the others, which are transparent. The DP and Gate are also shown in the T (left) and O (right) monomers. A model substrate of AcrB is also shown in CPK representation and colored gray. A black dotted arrow on the right picture indicates the direction of transport. (C) Same as in B, but viewed from the top.

The feasibility of the functional rotation mechanism at a molecular level, however, remains to be established. Indeed, while the need for concerted conformational changes in monomers of AcrB was demonstrated by several experiments (25, 26), no study addressed so far if and how substrate transport really occurs through the proposed mechanism. In fact, neither the molecular determinants nor the energetics of the process have been elucidated to date. Direct inspection of the functional rotation mechanism at an atomic level will ultimately provide a better understanding of how RND-type transporters work (possibly shedding light on rules governing polyspecific transport) and precious information for antibacterial drug discovery. In particular, the knowledge of the mechanistic details of the L<u>T</u>O→T<u>O</u>L step of the functional rotation (the underline indicates the monomer transporting the substrate; hereafter T→O) would represent a key milestone. Indeed, wherever recognition occurs, substrates should transit through the DP and the Gate in order to reach the Funnel domain of AcrB (17, 18, 20, 21).

Access to atomistic dynamics of complex molecular machines can be achieved nowadays by means of computer simulations, which represent an alternative and complementary approach to biochemical, biophysical, and structural experiments (27–34). As obtaining high-resolution structures of the complexes between RND transporters and known “good” substrates proved to be very challenging (12), it is particular important to develop computational protocols able to understand mechanisms such as those involved in the extrusion of substrates by the RND efflux pumps (35).

A few computational studies by us and other groups investigated in part the functional rotation mechanism in AcrB by using either enhanced-sampling all-atom molecular dynamics (MD) simulations (36–40) or a coarse-grained representation of the protein and its substrates (41, 42). While these studies provided the first insights into the link between conformational changes in the protein and translocation of substrates, they were limited in scope.Specifically, the coarse-grained approaches employed to study RND transporters used a single bead to represent each amino acid in the protein, thus they cannot dissect the role of different atomic-level interactions occurring along the translocation pathways of the substrates. In particular, they cannot address the role of water in the process, which is likely to be important for translocation as substrate diffusion channels are filled with solvent. Regarding the all-atom simulations performed to date: i) they were relatively short considering the size of the system under study, and ii) the conformational changes in the protein and the translocation of compounds were decoupled. These limitations hampered a quantitative understanding of how the former process drives the latter. Most importantly, no study evaluated (to the best of our knowledge) the energetics associated with the diffusion of compounds from the DP to the Funnel domain of AcrB during the T"O conformational change. Thus, we don't know how the functional rotation mechanism facilitates the diffusion of substrates of RND tranporters.

Prompted by these considerations, we developed a computational protocol based on multiple-bias MD simulations to characterize for the first time the key step T→O of the functional rotation mechanism, assessing its energetics and the role of the solvent in the process. The protocol was employed to study the translocation of doxorubicin (hereafter DOX, which until very recently was, together with minocycline, the only drug co-crystallized within the DP of AcrB (17, 23, 43)) from the DP to the Funnel domain *driven* by conformational changes occurring in AcrB.

Our results demonstrated the effectiveness of the functional rotation mechanism in facilitating smooth transport of substrates. The peristaltic motions occurring within internal channels of AcrB enabled a water layer soaking the internal surface of the translocation channel. This in turn permitted a fairly constant wetting of DOX during transport. The mediating action of water leveled off the free energy profile associated to the displacement of DOX. We speculate that the above mechanism can be generalized to rationalize the transport of substrates other than DOX, and that the mediating action of water is crucial for polyspecific transport by AcrB. Plausibly, other multi-drug transporters could exploit a very similar mechanism to perform their biological functions.

## Results and Discussion

In this section we first describe our computational protocol in brief. Then, we summarize and discuss our main results, comparing them with the available experimental data. We describe the structural, dynamical and energetic features of the functional rotation mechanism as seen in our simulations.

### Multi-bias MD simulations allow mimicking the functional rotation mechanism in silico

The conformational changes occurring in AcrB along the T→O step of the functional rotation were simulated by means of targeted MD (TMD) (44). This technique mimics structural rearrangements involving large protein regions by applying, to selected atoms (Cas here), a force proportional to their displacement from a linear path connecting the initial and final structures. DOX translocation from the DP to the Funnel domain within AcrB was induced by steered MD (SMD), in which one end of a virtual spring is attached to the molecule and the other end is pulled along a predefined direction (45, 46).

The novelty of our approach consists in efficiently coupling these two techniques in order to thoroughly mimic the functional rotation *in silico*, i.e. simulating substrate translocation *driven* by the structural changes of AcrB. In order to achieve this goal, we first addressed a key issue concerning the coupling times between protein conformational changes and substrate translocation. Indeed, different substrates could “respond” (i.e. unbind from the DP) with different timings to the conformational changes in the RND transporter, due e.g. to their different binding affinities, sizes, etc. Moreover we do not know a priori the details of such process even for a single substrate, including DOX.

To cope with this issue, we simulated different possible couplings between the two processes. Namely, we performed three simulations in which DOX was pulled from the DP towards the Funnel domain in 1 µs, while the T→O conformational transitions in AcrB were induced respectively within the first 0.1, 0.2, and 0.3 µs (Table S1). Hereafter, we refer to these simulations as *T_rot__10%T_pull_*, *T_rot__20%T_pull_* and *T_rot__30%T_pull_*, respectively. In addition, we performed a 1 µs long SMD simulation without concomitant induction of conformational changes in AcrB (referred to as *T_pull_1µs_*). Though not representative of any putative transport mechanism, the *T_pull_1µs_* simulation is very useful for comparative purposes.

Finally, to quantitatively address to what extent the functional rotation mechanism does promote the diffusion of substrates, we evaluated the free energy profile associated with translocation of DOX by means of umbrella sampling (US) simulations (47, 48) (Tables S1 and S2).

### The in silico substrate translocation mechanism is consistent with experimental data

First, we validated our computational protocol by assessing the consistency of the mechanism of substrate translocation *in silico* with the available experimental data.

Overall, the translocation of DOX as seen in our simulations turns out to be quite unaffected by the details of the computational setup, *as far as the conformational changes of the protein are mimicked concomitantly to the displacement of the substrate*. Importantly, the substrate is always transported through the putative Gate of AcrB (Figure 2A-B). More generally, during translocation DOX interacts with protein residues suggested experimentally (49) to be part of the extended DP and of the putative Gate (Figure 2C). Being an extension of the vast and malleable DP, the path toward the Gate enables consistent substrate rearrangement, as documented by the change in the orientation of DOX observed in all simulations (Figure S1). Previous experimental data demonstrated that DOX binds to the DP of AcrB assuming almost flipped orientations (17, 23) (Figure S2). Our results show that rotation of the substrate is possible also during its transport, at least within the channel leading from the upper part of the DP to the Gate. Importantly, such reorientation of DOX occurs at almost no cost (see below). The simulated translocation process is thus compatible with several experimental findings.

**Figure 2.**
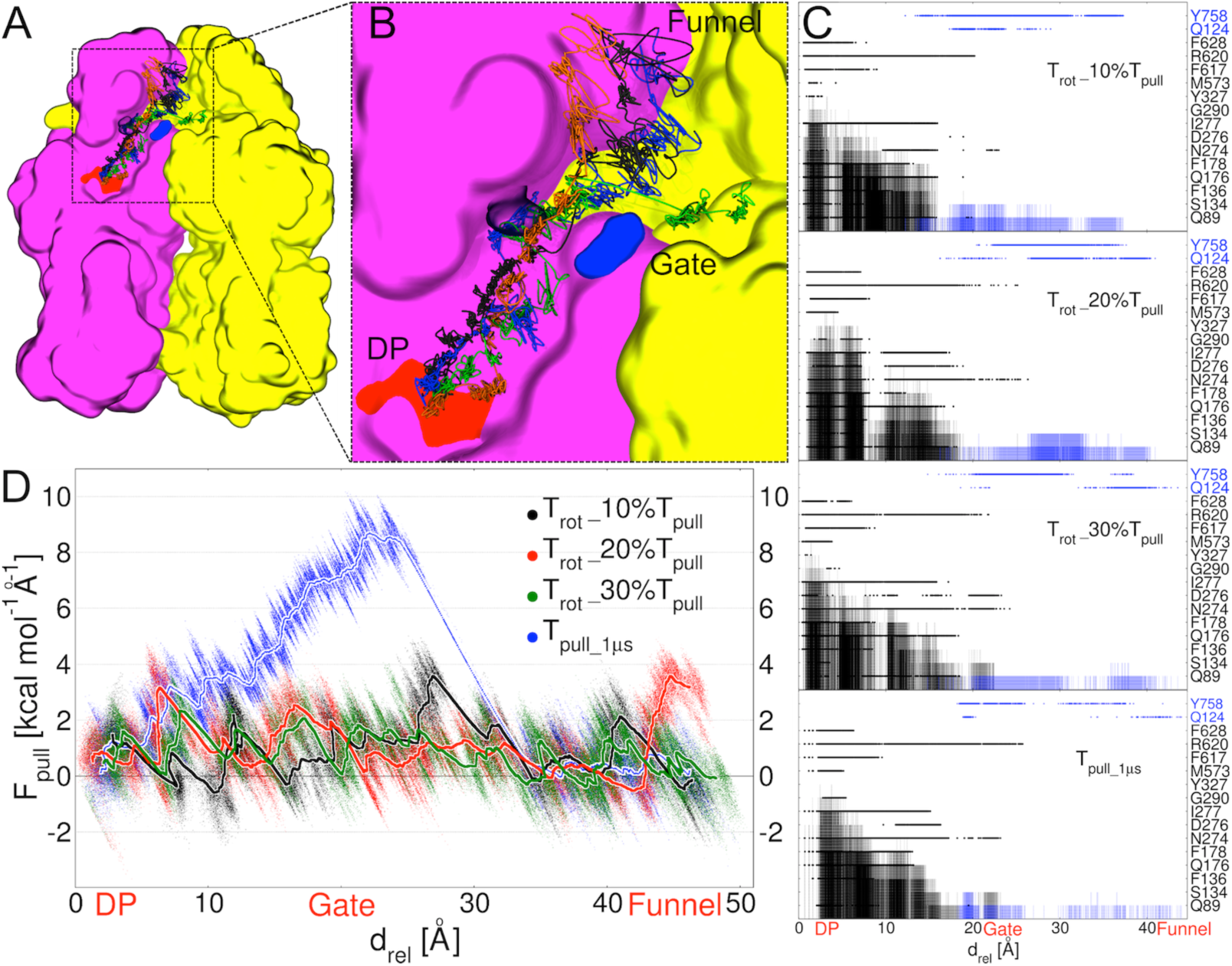
Translocation of DOX during the T→O step of the functional rotation in AcrB. (A) Overall view of translocation pathways from the DP to the Gate as seen in multi-bias MD simulations labeled *T_rot__10%T_pull_*, *T_rot__20%T_pull_* and *T_rot__30%T_pull_* and in the SMD simulation *T_pull_1µs_*. For clarity, only monomers T and L of AcrB are shown in magenta and yellow surfaces, respectively. In addition, some regions of the monomer T were removed to allow visualizing the transport pathways. Part of the DP within monomer T is colored red. Residues Q124 and Y758 lining the Gate are shown as solid and transparent blue surfaces, respectively. Black, red, green and blue solid lines indicate pathways of DOX in *T_rot__10%T_pull_*, *T_rot__20%T_pull_*, *T_rot__30%T_pull_* and *T_pull_1µs_*, respectively. (B) Magnification of the region within the square dotted box in A. (C) Plots of the per-residue (dots) and cumulative (semi-transparent bars) contacts between DOX and residues identified as those lining the substrate transport pathway in AcrB (49). The x-axis reports the displacement *d_rel_* of DOX with respect to its initial position within the DP. Black and blue labels indicate residues lining the DP and the Gate, respectively. A contact was recorded at each step if the minimum distance of DOX from any residue was lower than 3 Å. (D) Profiles of *F_pull_* extracted from *T_rot__10%T_pull_*, *T_rot__20%T_pull_*, *T_rot__30%T_pull_* and *T_pull_1µs_*. Tiny points and thick lines indicate forces calculated every 2000 steps of MD simulation and running averages over 5000 points, respectively.

### The functional rotation facilitates diffusion of substrates by lowering conformational strain and allowing for their continuous wetting within the transport channel

The overall agreement among the main results obtained from *T_rot__1/2/30%T_pull_* simulations discussed above is congruous with the resemblance of the profiles of the pulling force *F_pull_* applied to induce the translocation of DOX (Figure 2D). Note that all profiles are relatively smooth and do not feature large values of *F_pull_*, which would be associated with transport bottlenecks. In particular, there is no evidence for high values of *F_pull_* near the Gate. On the contrary, a prominent peak of almost 10 kcal×mol-1 Å^-1^ in *F_pull_* appears near the Gate when the translocation of DOX is mimicked in the complete absence of conformational changes in AcrB, i.e., for *T_pull_1µs_* (Figure 2D). Thus, inducing the T→O conformational change in the first part of the simulation facilitates substrate translocation.

In order to shed light on the microscopic determinants of the process, we performed a detailed analysis of the interactions involving DOX, the protein and the solvent. As expected, in the absence of the T → O conformational change the passage of DOX through the Gate induces a clear structural strain in the substrate (Figure 3). More importantly, the opening of the Gate following the T →O transition enables a fairly structured hydration of almost the entire internal surface of the translocation channel leading from DP to the Funnel domain (Figure 4). This is particularly evident from the plot of the spatial distribution function (SDF) of water molecules (Figure 4A), which provides a picture of the order in liquid water and reveals specific details of its local structure (50). Note that structured hydration of the translocation channel is compatible with the relatively hydrophilic character of the channel connecting the DP to the Funnel domain (39).

**Figure 3.**
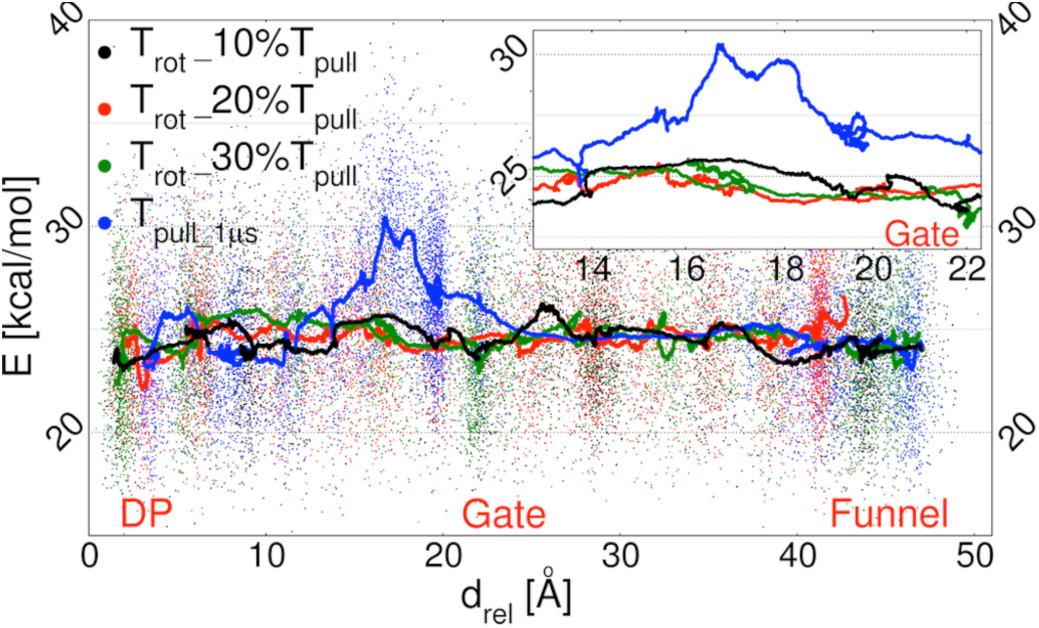
Conformational strain experienced by DOX during transport. Plots of the internal energy of DOX (calculated from the dihedral energy terms of the force field) calculated from simulations *T_rot__10%T_pull_*, *T_rot__20%T_pull_*, *T_rot__30%T_pull_* and *T_pull_1µs_*. Tiny points and lines indicate forces calculated every 2000 steps of the MD simulation and running averages over 5000 points, respectively. The inset shows the region of higher discrepancy between the profile from *T_pull_1µs_* and all the others.

**Figure 4.**
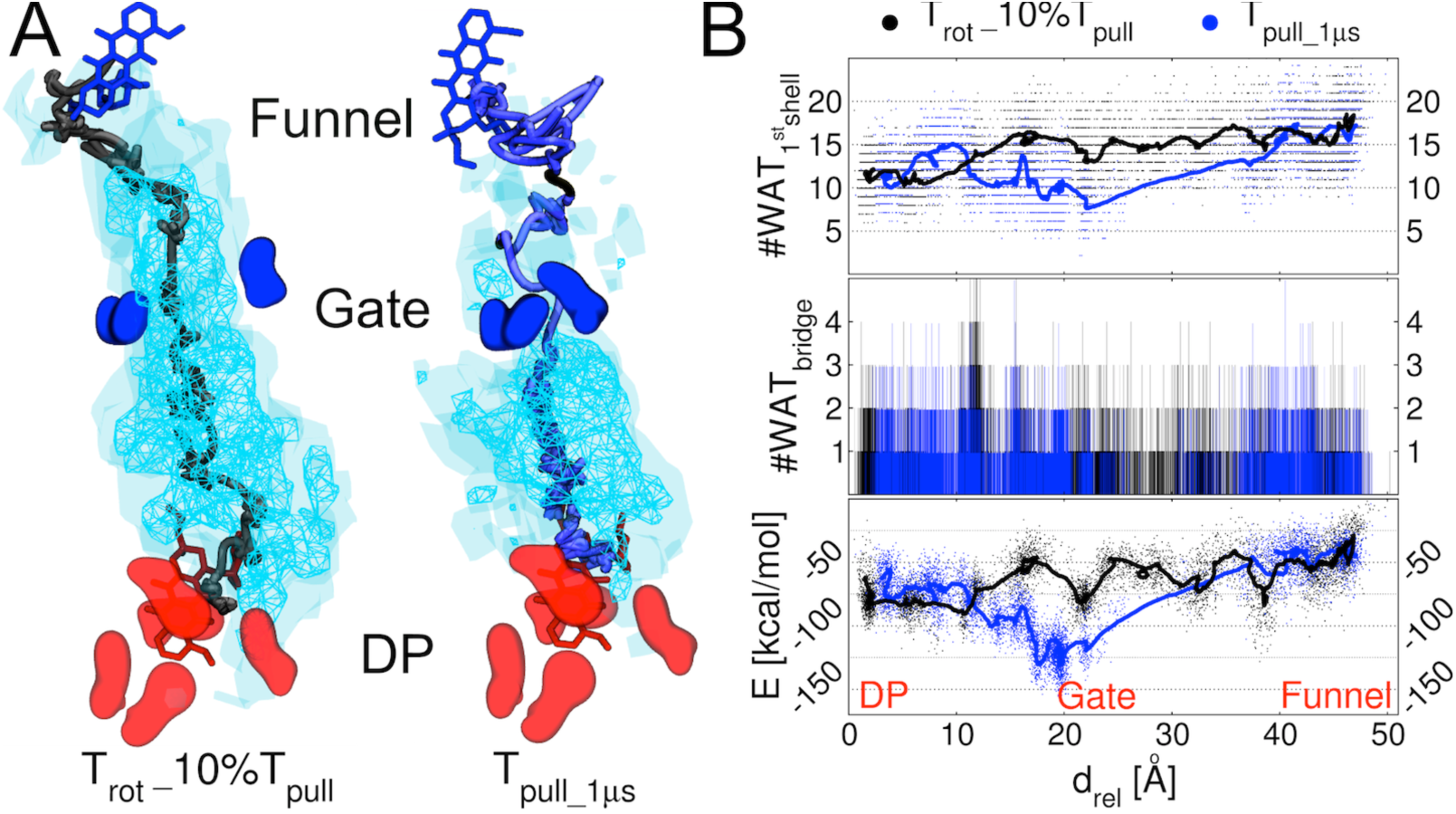
The functional rotation mechanism enables continuous hydration of the transport channel and of the substrate. (A) SDF isosurfaces of water oxygen atoms within the transport channel leading from the DP to the Funnel domain. Left and right pictures refer to simulations *T_rot__10%T_pull_* and *T_pull_1µs_*, respectively. The positions of DOX at the beginning and at the end of the simulations are shown as red and blue sticks, respectively. The pathways traced by the center of mass of the drug (sampled every 2.5 ns) in *T_rot__10%T_pull_* and *T_pull_1µs_* are displayed as dark grey and light blue tubes respectively. SDF surfaces corresponding to isovalues of 5 and 1 (with respect to the average value in bulk water) are shown as cyan nets and transparent surfaces, respectively. (B) DOX hydration properties and interaction with AcrB during transport in *T_rot__10%T_pull_* and *T_pull_1µs_*. Upper graph: number of water molecules within the first hydration shell of DOX as a function of drel. Middle graph: number of water-bridged interactions between DOX and the protein. Lower graph: profiles of AcrB-DOX interaction energy calculated from the force-field terms representing electrostatics and van der Waals interactions. Tiny points and lines in the upper and lower graphs indicate respectively values calculated every 2000 steps and running averages over 5000 points. See Figure S3 for a comparison including also *T_rot__20%T_pull_* and *T_rot__30%T_pull_*.

By comparing the hydration properties of the translocation channel and of the substrate in *T_pull__10%T_rot_* and *T_pull_1µs_* simulations, it is clear that this phenomenon is a feature of the functional rotation mechanism (Figure 4). As a consequence of the reduced screening effect of waters in *T_pull_1µs_*, a negative peak appeared in the corresponding DOX-AcrB interaction energy profile (Figure 4B and Figure S3). The presence of a roughly continuous layer of waters within the translocation channel enables a fairly constant wetting of the substrate, as well as several water-mediated interactions with the protein (Figure 4B).

### Substrate translocation occurs over a smooth free energy profile

We calculated the free energy profile associated with the transport of DOX from the DP to the rear of the Gate (Figure 5), that is the part of the translocation process occurring within the AcrB channel (corresponding to a change of about 35 Å in the relative displacement from the DP, *d_rel_*). The profile is relatively smooth, with barriers lower than 5 kcal×mol-1. Furthermore, the affinities of DOX to the DP and the Gate are virtually identical (Figure 5A); thus, the probability to find the substrate near the second site increases in response to conformational cycling in AcrB (see next section for biological implications).

We noticed that the orientation of DOX along its translocation path changes near the Gate (inset in Figure 5B, and Figure S1). Therefore, we included an angular variable *a_DOX/DP-Funnel_* representing DOX rotation (see Materials and Methods) to estimate a 2D free energy profile in the corresponding region of the translocation process (Figure 5C). Importantly, the addition of *a_DOX/DP-Funnel_* had a very small impact on the results discussed above. Indeed, both the free energy difference found between the states representing almost flipped orientations of DOX (labeled 3 and 4 in Figure 5A) and the barrier associated with the rotation remain vanishingly small.

**Figure 5.**
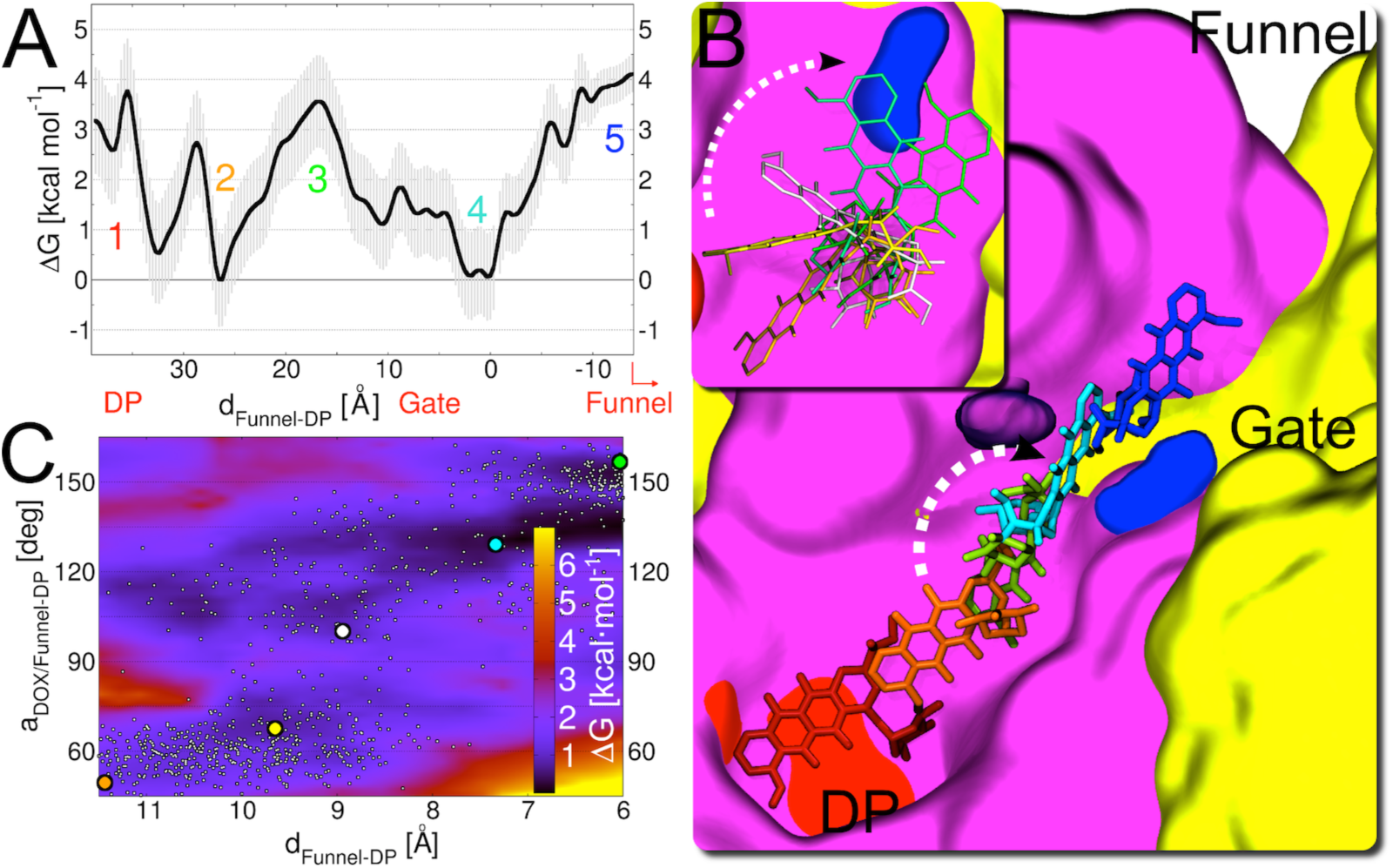
Energetics of DOX transport within AcrB. (A) 1D free energy profile associated to transport of DOX from the DP to the rear of the Gate along the T→O step of the functional rotation. The profile of ΔΔG (kcal×mol^-1^) referred to the binding free energy within the DP is reported as a function of the pseudo reaction coordinate *d_Funnel-DP_*, defined as the difference in the distance of DOX from the center of the Funnel domain and from the center of the DP (see Materials and Methods). Colored numbers in the graph identify arbitrary stages of the transport process, for which the conformation of DOX is shown in B with the same color code. (B) Representative conformations of DOX along the translocation pathway as seen in the *T_rot__10%T_pull_* simulation. DOX conformations are shown as sticks of different colors, representing different stages of the process and corresponding to positions identified by numbers 1 to 5 in A. The protein, the DP and the Gate are represented as in Figure 2A. Inset: detailed view of the rotation of DOX near the Gate. Selected conformations are shown as thin sticks colored as the large points in C. (C) 2D free energy profile as a function of *d_Funnel-DP_* and of the angle *a_DOX/DP-Funnel_*, calculated in the region where the change in orientation of DOX occurs (see Figure S1). Colored points indicate conformations of DOX extracted from the *T_rot__10%T_pull_* simulation and featuring different orientations, and shown with the same color code in the inset of B.

Clearly, a proper comparison of our results with experiments is flawed by several factors, including the lack of a free energy profile associated with substrate translocation through the whole AcrABZ-TolC efflux pump. However, it is worth noticing the absence in our profile of high free energy barriers, which would correspond to efflux kinetics inconsistent with the typical extrusion times of substrates by RND efflux pumps (up to 10^3^ compounds per pump per second depending on the substrate (51–53)).

### Biological implications

In this section we discuss further the possible implications of our findings in relation to the current view of how RND efflux pumps work.

#### AcrB substrates can adopt multiple binding modes also outside the DP

We showed that DOX assumes different orientations during transport. While such rotation can be facilitated by the increased hydration of DOX upon detachment from the DP (Figure 4B), we cannot exclude the presence of multifunctional sites (that is, sites able to recognize various types of functional groups, from hydrophobic to polar and charged) (54) within the channel leading from this site to the Gate. Note that the presence of such multifunctional sites has been demonstrated within the DP (54), and is coherent with the possibility for DOX to adopt (at least) two different orientations within this site (17, 23). A new possibility arising from our findings is that multidrug recognition and transport are not restricted to one or more binding sites (e.g. the DP), but rather dictated by the physico-chemical properties of the *entire* substrate translocation pathway through AcrB. These findings are compatible with polyspecific transport by AcrB.

#### A “one stroke - one drug” mechanism of substrate expulsion is not necessary for AcrB

As a result of the T → O conformational change in AcrB, the interaction strength of DOX with the Gate and with the DP are comparable (Figure 5A). Thus, AcrB could contribute to efflux by “just” favoring, through a functional rotation mechanism, the accumulation of substrates in the central region of the upper Funnel domain, beyond the Gate. This should create a concentration gradient driving translocation of a pile of compounds through the AcrA/TolC channel, as hypothesized earlier (24). Such a mechanism may explain how certain substrates (e.g. aminoacyl-β-naphthylamides (55)) are pumped out at rates far exceeding those expected for the common transporters, suggesting that many substrate molecules might be pushed out in one stroke.

The largely prevalent hydrophilic character identified in the internal surfaces of MexA and OprM (respectively homologous of AcrA and TolC in *Pseudomonas aeruginosa*) further corroborates our hypothesis (56). Moreover, an analysis of the hydrophilic character of the internal surfaces of AcrA and TolC in the recently published structure of the complete AcrABZ-TolC assembly (57) confirmed these findings (Figure S4). Therefore, it is plausible that upon crossing the Gate, the substrate will wander within an environment that would drive it out (56).

#### Transport rate bottleneck is not due to diffusion of substrates within AcrB

The smooth free energy profile associated with the translocation of DOX implies that the bottleneck in terms of rate of transport comes from the concerted conformational changes occurring in AcrB. This hypothesis is in line with the current understanding of how many active transporters work (58), and is supported by the comparison of our data with experimental studies reporting AcrB efflux rates of several compounds (51, 52, 59). For instance, the values of the turnover number *kcat* estimated by Nikaido and co-workers (51, 52) for the efflux of cephalosporins and penicillins range from ∼10 s^-1^ to ∼10^3^ s^-1^. Using simple arguments from transition state theory to get an approximate value of the effective free energy barrier Δ*DG‡* that would be compatible with this rate, we obtained ∼13 kcal.mol^-1^, which is well above the values found here. Interestingly, the effective free energy barrier calculated for the translocation of DOX through the TolC channel amounted to almost 10 kcal·mol^-1^ (60), a value similar to that extrapolated from experimental data. These findings further highlight the key effect of the conformational changes in AcrB on substrate diffusion, and support our statement about efflux rate bottlenecks.

#### Water is key for polyspecific transport by AcrB

According to our results, the relief of steric hindrance and the formation of a continuous layer of structured waters crucially facilitate substrate transport inside AcrB. The mechanism we propose would: *i)* match with the increasing ratio of hydrophilic over hydrophobic residues along the channel leading from the DP to the Funnel domain (39); *ii)* be compatible with the many structural waters found within internal surfaces of AcrB in the highest-resolution crystal structure reported to date (PDB ID 4DX5 (23)).

Moreover, it is likely that this very general mechanism would facilitate diffusion of several chemically unrelated substrates dissociating from the DP, thus enabling polyspecific transport. Indeed, by shielding potentially (too) strong interactions between chemical groups of compounds and AcrB, water would in part “hide” chemical differences among diverse substrates. Therefore, translocation of neutral, zwitterionic, anionic and cationic compounds could occur along a similar path and with comparable overall costs. Although verifying such a hypothesis with other compounds would be very computationally demanding, we point out that among the drugs co-crystallized so far within the DP of AcrB (including minocycline (17, 23) and puromycin (43)), DOX features overall the largest molecular mass, van der Waals volume and minimal projection area (see e.g. data at https://www.dsf.unica.it/translocation/db (61)). Moreover, it is as soluble as the other substrates (all have high solubility in water, the values of intrinsic logS being −3.6, −3.2 and −2.3 for DOX, puromycin and minocycline, respectively according to Chemicalize - <u>https://chemicalize.com</u>). Therefore, it is plausible to expect that smaller and similarly soluble compounds than DOX could be transported via the mechanism described above. These considerations strongly suggest that water mediates the transport of (at least) low-molecular mass substrates recognized at the DP.

#### Recognition and transport of substrates in AcrB

On the basis of our findings and previous literature, the mechanism by which AcrB and its homologs proteins recognize and transport their substrates would be as follows:

- Recognition occurs via interaction of substrates with one among the multiple binding sites present in AcrB, each endowed with a few multifunctional sites (17, 20, 21, 23).
- Concerning the DP, the interaction of the substrate with this site triggers the T→O conformational changes in the protein, which decreases the affinity of the compound.
- Upon unbinding from the DP, the substrate will find itself in a relatively hydrated environment favoring smooth diffusion towards the Gate or, at least, disfavoring specific interactions that could hinder transport. This hypothesis matches with the physico-chemical traits of typical AcrB substrates, whose unique common feature is some degree of lipophilicity (5, 16). Furthermore, the latter scenario is compatible with the transport of a pile of compounds through repeated conformational cycling of the pump (24).

Note that, according to the mechanism described above, the distinction between a substrate and an inhibitor should be in the way they bind to the same region of AcrB (binding strength and/or position), which should induce different rearrangements of the pump, as already suggested by several studies (43, 62–64).

## Conclusions and Perspectives

In this work we validated a novel computational protocol to mimic *in silico* the key step (T→O) of the functional rotation mechanism by which RND transporters such as AcrB are believed to export their substrates. To the best of our knowledge, this is the first computational study: *i)* addressing the coupling between the conformational transitions occurring in RND-type transporters and the translocation of its substrates, and *ii)* providing an estimation of the free energy profile associated with the key step of the process, that is the transport of a substrate from the Distal Pocket to the Funnel domain. Thanks to this unprecedented computational effort we characterized the molecular determinants of substrate translocation caused by peristaltic-like motions occurring within internal channels of AcrB. Using doxorubicin as a probe we showed how these structural changes favor substrate transport along a path that is fully compatible with that proposed on the basis of X-ray data and whole cells assays. Moreover, we propose a rationale for the polyspecific transport by the RND-type multidrug efflux pump AcrB, in which water molecules play a key role. Clearly, water-mediated transport could be a general feature of the multidrug transport mechanism.

Accurate computational protocols such as the one used here represent a valid and highly informative strategy to understand the molecular mechanisms of recognition and transport by RND proteins. Moreover, given the robustness of the methodology with respect to implementation details, we are confident that it can be successfully applied to study the transport of other substrates by AcrB and homologous proteins, and easily adapted to investigate complex processes in other biological systems.

## Materials and Methods

In the following we describe in detail the system we studied and the computational protocol we employed. We also discuss the possible limitations of our approach. An extensive validation of the methodology and a comparison of our results with previously published computational work are reported in the Supplementary Information.

### Simulated System

The system under study has been described in detail earlier (36). The starting structure consisted of a molecule of DOX bound to the DP of monomer T of the asymmetric homotrimeric AcrB, and was taken from the equilibrium dynamics reported in (36). The protein was embedded in a 1-Palmitoyl-2-Oleoyl-PhosphatidylEthanolamine (POPE) membrane bilayer model, and the whole system was solvated with a 0.15 M aqueous KCl solution. The total number of atoms in the system was ∼450,000.

The parameters of DOX (freely available at <u>http://www.dsf.unica.it/translocation/db</u>(61)) were taken from the GAFF force field (65) or generated using the tools of the AMBER 14 package (66). In particular, atomic restrained electrostatic potential (RESP) charges were derived using antechamber, after a structural optimization performed with Gaussian09 (67). The force field for POPE molecules was taken from(36). The AMBER force field parm14SB (68) was used for the protein, the TIP3P (69) model was employed for water, and the parameters for the ions were taken from ref. (70).

### Computational protocol

In order to mimic DOX translocation from the DP to the Funnel domain in response to the T→O conformational transition of AcrB, we devised an original computational protocol that couples two well-established methods to enhance the sampling of biological processes. Namely, we mimicked the conformational change of the protein via TMD simulations (44) *while* pulling the substrate towards its putative path through the Gate by means of SMD simulations (45, 46). To the best of our knowledge, these two computational methodologies were never combined to date. The resulting DOX translocation pathway was discretized into several snapshots used as starting conformations to perform 1D and 2D US MD simulations (47, 48). Finally, the weighted histogram analysis method (WHAM) (71) was used to estimate the free energy profile associated to the transport of DOX from the DP to the rear of the Gate. Simulations were analyzed with in-house *tcl, bash* and *perl* scripts, with the *cpptraj* tool of the AMBER 14 package (66) and with tools of the GROMACS 5.1.4 package (72). Figures were prepared using xmgrace, gnuplot and VMD 1.9.2 (73).

#### Targeted and steered MD simulations

These simulations (Table S1) were performed using the NAMD 2.9 package (74). A time step of 1.5 fs was used to integrate the equations of motion. Periodic boundary conditions were employed, and electrostatic interactions were treated using the particle-mesh Ewald method, with a real space cutoff of 12 Å and a grid spacing of 1 Å per grid point in each dimension. The van der Waals energies were calculated using a smooth cutoff (switching radius 10 Å, cutoff radius 12 Å). MD simulations were performed in the NPT ensemble. The temperature was maintained around 310 K by applying Langevin forces to all heavy atoms, with a damping constant of 5 ps-1. The pressure was kept at 1 atm using the Nosé-Hoover Langevin piston pressure control with default parameters.

TMD (44) simulations allowed mimicking the conformational transitions between the two known conformational states (L<u>T</u>O and T<u>O</u>L) of AcrB. It was recently shown that TMD simulations produce reliable transition paths as compared to other more refined techniques (75). To prevent any steric hindrance induced on the T monomer by the neighbors, we targeted all of them toward their next state along the functional rotation cycle. Furthermore, to allow for the largest conformational freedom of the protein along the pathway traced by the targeting restraints, these were applied only to the Cα atoms of AcrB. The force constant was set to 1 kcal·mol^-1^·Å^-2^ for each carbon atom, much lower than that employed in earlier studies (36, 39).

Regarding SMD simulations (45, 46), a relatively low force constant (1 kcal·mol^-1^·Å^-2^) was applied to the center of mass of the non-hydrogenous atoms of the substrate. This choice allowed the molecule deviating from a straight pathway and optimizing interactions with the surrounding environment during transport.

#### Umbrella sampling simulations

To estimate the energetics associated with the translocation of DOX we performed extensive 1D and 2D US simulations (47, 48) along the pathway from the DP to the rear of the Gate (corresponding to a displacement of about 35 Å). The pseudo reaction coordinate used for the evaluation of the free energy profile is different from that used to plot data from the targeted/steered MD simulations. Indeed, in the former simulations we calculated the force as a function of *d_rel_*, the distance traveled by the center of mass of the substrate with respect to its initial position. The free energy profile was plotted instead as a function of *d_Funnel-DP_*, defined as the difference *d_DOX-Funnel_ - d_DOX-DP_* between the distance of the center of mass of DOX from the centers of mass of the Funnel domain and of the DP. This choice provides a finer grid for the evaluation of the free energy profile. The path of DOX was discretized into 35 windows covering a *d_Funnel-DP_* range of about 50 Å and placed at 1.5 Å from each other, starting at *d_Funnel-DP_* = 38.0 Å (Table S2). The force constant *k_d_* used to restrain the sampling of DOX within each window along *d_Funnel-DP_* was set to 5 kcal·mol^-1^·Å^-2^. 4 additional windows were added to obtain uniform sampling across the pathway, namely at values of *d_Funnel-DP_* of −9.2, −1.8, 28.3 and 34.3 Å.

Since the orientation of DOX changed near the Gate (namely at values of *d_Funnel-DP_* between ∼5 Å and ∼12 Å; Figure 5 and Figure S1), a single reaction coordinate could be insufficient to evaluate the free energy profile correctly. Therefore, a 2D free energy surface was evaluated in that region by introducing the additional angular variable *a_DOX/DP-Funnel_*. This is defined as the angle between the main axis of DOX (approximately identified by the line connecting two atoms of the tetracyclic body of the molecule) and the line connecting the centers of mass of the DP and of the Funnel domain (Figure S5). A total of 28 simulations, each of 25 ns in length, were performed on a grid defined by points of coordinates (*d_Funnel-DP_, a_DOX/DP-Funnel_*) = (*{6.5, 8, 9.5, 11} Å, {60, 75, 90, 105, 120, 135, 150} °*). The same value of *k_d_* as in the 1D US simulations was used to restrain the sampling along *d_Funnel-DP_*, while a force constant *ka* of 120 kcal·mol-1·rad-2 was used to restrain the orientation of the substrate.

The values of *d_Funnel-DP_* and *a_DOX/DP-Funnel_* were saved every 2 ps (corresponding to 1333 simulation steps). The WHAM method (71) as implemented in the *g_wham* tool of GROMACS was used to extract the free energy profiles and surfaces, using a tolerance of 10^-6^ for the convergence of the probability. Simple Bayesian bootstrapping was utilized to estimate the statistical sampling errors using 500 randomly chosen data sets with the same data size.

The RMSD of the complexes with respect to their initial conformation revealed minor structural changes in all windows (Figure S6A). A fairly flat profile was indeed reached after ∼20 ns. In addition, the orientation of DOX did not change significantly in any of the 1D US windows (Figure S6B). The relatively good stability of the systems was mirrored in the fairly good convergence of the free energy profile (Figure S6C). The free energy profile (surface) reported in Figure 6A (B) was estimated over the last 12.5 (6.25) ns of the simulation.

### Limitations of our approach

Our methodology is based on all-atom classical MD simulations with predefined protonation states of all molecules. As such, it neglects the coupling between conformational changes occurring in the periplasmic region of AcrB and the flux of protons across the TM domain. However, this limitation will hardly affect the outcome, since the translocation of DOX occurs fully within the periplasmic domain. Furthermore, we are mimicking exactly the process (i.e. the L<u>T</u>O → T<u>O</u>L conformational change) induced by the change in the protonation states of key residues within the TM region.

Another limitation of our approach consists in neglecting the AcrB partners forming the full AcrABZ-TolC assembly (43, 57, 76, 77). In particular, the extrusion process could be affected by the interaction between AcrB and AcrA, which could alter e.g. the flexibility and the hydration properties of the upper part of the Funnel domain. However, we believe that restricting our study to AcrB does not constitute a major drawback, as the translocation of DOX simulated here occurs mainly within internal AcrB channels.

## Acknowledgments

The research leading to the results discussed here was partly conducted as part of the Translocation Consortium (http://www.translocation.eu) and has received support from the Innovative Medicines Initiative Joint Undertaking under Grant Agreement no. 115525, resources that are composed of financial contribution from the European Union's Seventh Framework Programme (FP7/2007-2013) and EFPIA companies in kind contribution. VKR is a Marie Skłodowska-Curie fellow within the “Translocation” Network, project no. 607694. This research used the Savio computational cluster resource provided by the Berkeley Research Computing program at the University of California, Berkeley (supported by the UC Berkeley Chancellor, Vice Chancellor for Research, and Chief Information Officer). We thank H. Nikaido (University of California at Berkeley, USA), K.M. Pos (Goethe University, Frankfurt am Main, Germany), H. Zgurskaya (University of Oklahoma, Norman, OK, USA), M. Picard (CNRS/Université Paris-Diderot, Paris, France), E. Bibi (Weizmann Institute of Science, Rehovot, Israel), J. Blair and L. Piddock (University of Birmingham, UK), C. Melis (University of Cagliari, Italy), F. Pietrucci (Université Pierre et Marie Curie, Paris, France) and A. Kranjc for the critical reading of the manuscript.

## Author Contributions

A.V.V, U.K. and P.R. conceived and designed research. A.V.V. performed research. A.V.V. and P.R. analyzed the data. A.V.V., V.K.R., I.M. and G.M. contributed materials and analysis tools. All authors wrote the manuscript.

